# Germ-cell restricted antigens are presented on HLA-I in premalignant colorectal lesions

**DOI:** 10.64898/2026.11.25.768254

**Authors:** Shubham Gupta, Suvi Karumo, Ruben Shrestha, Hayan Lee, Edward D. Esplin, Molly Miranda, Hoang Tran, Joanna Bi, Rozelle Laquindabaum, Emma Monte, Shannon M. White, Ruiqi Jian, Xuhuai Ji, Uri Ladabaum, Otto K. Kari, James M Ford, Dansen Hornburg, Michael P. Snyder

## Abstract

Tumour-restricted antigens displayed by HLA class I proteins are potential targets for T cell recognition, but whether they are present before a lesion becomes invasive is unknown. Here we applied high-sensitivity HLA-I immunopeptidomics to 105 individual colorectal lesions and normal mucosa from 24 donors with hereditary polyposis. We identify peptides from germ-cell-restricted genes presented in premalignant lesions, confirmed against synthetic standards, and absent from reference immunopeptidomes of benign colon. These source genes are hypomethylated in polyps relative to normal mucosa. The same pattern is seen in colorectal tumours, and demethylating agents induce their expression in cell lines. Peptides were presented from transcripts at abundances well below conventional thresholds for target nomination, demonstrating the relevance of direct immunopeptidome readout with mass spectrometry. Several source genes were silent in medullary thymic epithelium, and dextramer-sorted T cells against one such peptide were dominated by a small number of clonotypes. An antigenic landscape therefore emerges in colorectal tissue before malignant transformation.

## Introduction

HLA class I molecules display intracellularly derived peptides at the cell surface, providing T cells with a direct readout of cellular state. HLA immunopeptidomics uses mass spectrometry to identify these presented peptides following isolation of peptide-HLA complexes, enabling direct measurement of the antigenic landscape without relying on transcript abundance or binding prediction [1]. This is particularly important in cancer, where expression alone cannot establish whether an antigen is processed, loaded onto HLA, and displayed. Advances in instrument sensitivity, including trapped ion-mobility, now enable deep immunopeptidome profiling from limited tissue material, bringing small specimens such as premalignant lesions within reach [2,3].

Colorectal immunopeptidomes have been characterised in both microsatellite stable and microsatellite unstable tumours [4–7], and the HLA Ligand Atlas provides a reference peptidome of benign colon from autopsy donors [8], against which specimens lacking matched normal tissue can be compared. How this repertoire changes in precancerous lesions, however, has not been examined.

In precancerous polyps, low tumour mutational burden [9] makes neoantigens scarce and non-mutated self-antigens the more plausible source of tumour-restricted peptides. These lesions are also where loss of the HLA-I complex first appears [10], suggesting early immune escape. Such antigens are likely to derive from genes whose expression is normally confined to germ cells and other immune-privileged sites, commonly called cancer-testis antigens (CTAs), and silenced in somatic tissue by DNA methylation [11–13]. Demethylating agents can induce their presentation on HLA-I in cell lines [14,15]. Whether CpG demethylation derepresses these loci in premalignant human tissue, and whether their peptides reach the cell surface before a lesion becomes invasive, remains to be seen.

CTAs such as MAGE family members, NY-ESO-1 [12] and PRAME [13,16] established the principle that immune-privileged genes can supply antigens absent from healthy somatic cells (GTEx) [17]. Interventional trials built on such peptides have nonetheless produced limited benefit [18–20], which makes it important to know whether the immune repertoire against tumour-restricted antigens is already tolerized [21].

We studied colorectal polyps and normal mucosa from familial adenomatous polyposis (FAP) patients carrying germline APC mutations. Because somatic APC loss also initiates the majority of sporadic colorectal cancers (85%), lesions in these patients model the earliest events of the adenoma-carcinoma sequence [22]. Individuals with FAP undergo surveillance colonoscopy starting as early adolescence and prophylactic colectomy as young adults, which provides access to polyps with matched normal mucosa that is rarely sampled in sporadic disease. We performed ultra high sensitivity HLA-I immunopeptidomics on these low-mass specimens, and interpreted the results against transcriptional [9] and CpG methylation data [23]. We show that DNA hypomethylation in premalignant lesions is accompanied by derepression of germ-cell-restricted genes whose products are processed and presented on HLA-I. Analysis of thymic expression and dextramer-sorted T cell repertoires further suggests that immune recognition of selected antigens may be retained. Together these findings reveal an antigenic landscape that emerges before malignant transformation.

## Results

### dda-PASEF and dia-PASEF immunopeptidomics of individual precancerous lesions

We isolated HLA-I–bound peptides from 84 colorectal polyps, 16 normal and 5 malignant tumor samples and analyzed them using high-sensitivity mass spectrometry (Figure 1a, Methods). Although polyps were small (∼20 mg), each of the 105 specimens was processed individually rather than pooled, allowing per-lesion peptidomes to be resolved [24]. Peptides from specimens were split between data-dependent acquisition (dda-PASEF) to build the spectral library (n = 114) and data-independent acquisition (dia-PASEF) for quantification (n = 89). Specimens were derived from 24 donors which included 12 with familial adenomatous polyposis, 3 with MUTYH-associated polyposis (MAP), 2 normal autopsy, one sporadic cancer patient, and 6 donors contributing a single case each (Figure 1b, supplementary Figure 1a).

**Figure 1.**
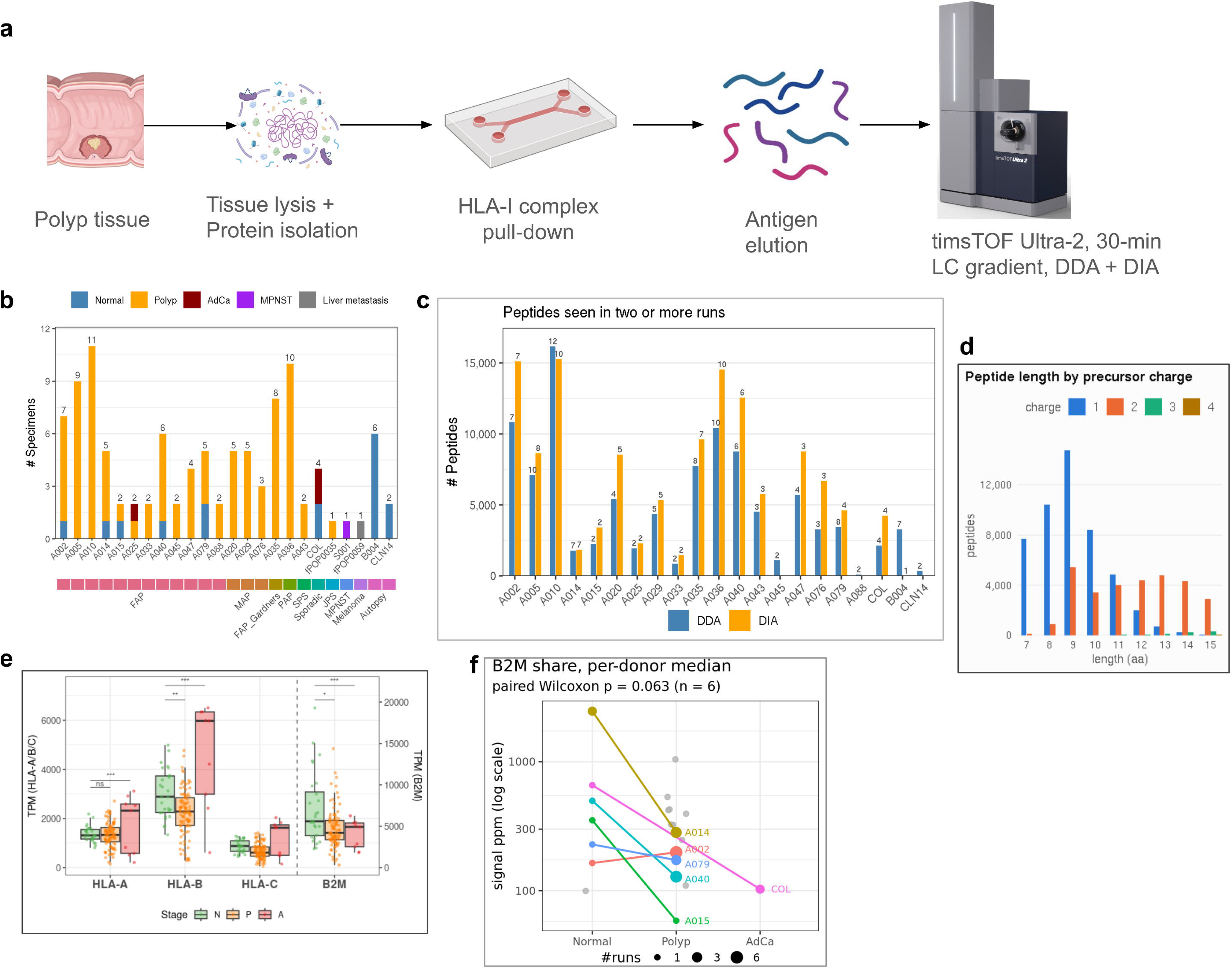
HLA-I Immunopeptidomics workflow and cohort. **a**) Schematic of the experimental workflow. Specimens were lysed and HLA-I complexes were captured on PeptiCHIP cartridges, and eluted peptides acquired on a Bruker timsTOF Ultra 2. **b**) Number of specimens used per donor, coloured by tissue type. The disease status of each donor is indicated by the annotation bar below. There were 105 specimens from 24 donors. **c**) Peptides detected in two or more runs per donor, shown separately for DDA (blue) and DIA (orange). Numbers above bars give the number of samples contributing to each acquisition mode. **d**) Peptide length distribution by precursor charge state for DIA-quantified peptides. **e)** Transcript abundance of HLA-I genes on the left axis and Beta 2 microglobulin (β2M) on the right axis, across disease stages -N: Normal, P: Polyp, A: Adenocarcinoma. **f)** β2M abundance expressed as its share of the total immunopeptidome signal (signal ppm, log scale) across normal, polyp and adenocarcinoma (AdCa) tissue. Data comprise 83 runs from 17 patients; the 6 patients with more than one tissue type sampled are shown as coloured points joined by lines, and patients with a single tissue are shown in grey. β2M signal declines from normal to polyp in most paired patients. AdCa: adenocarcinoma; dda-PASEF: data-dependent acquisition with parallel accumulation-serial fragmentation; dia-PASEF: data-independent acquisition with parallel accumulation-serial fragmentation; FAP: familial adenomatous polyposis; JPS: juvenile polyposis syndrome; MAP: MUTYH-associated polyposis; Melanoma: blue-nevus melanoma; MPNST: malignant peripheral nerve sheath tumour; PAP: polymerase proofreading-associated polyposis; SPS: serrated polyposis syndrome

From dda-PASEF (data-dependent acquisition-Parallel Accumulation Serial Fragmentation), we identified 138,979 peptides, 74.7% of which fell within the canonical 8-12 amino-acid (aa) length range. dia-PASEF resulted in 76,420 peptides, with 73% in the canonical length range. Most DDA identifications were singletons (supplementary Figure 1b,c), whereas dia-PASEF recovered 39.1% of peptides in more than one specimen per donor (Figure 1c). Among dia-PASEF peptides of canonical length, 71.1% carried a +1 charge, with higher charge states becoming more common as peptide length increased (Figure 1d). Our data acquisition strategy followed Phulphagar et al. (2023), which recommends including +1 precursors and lower collision energies for immunopeptidomics than conventional proteomics, where +2 precursors and higher collision energy predominate. A median of 13,237 peptides was identified per donor and 4,908 per specimen (range: 213 -14,826). We next quantified how often each peptide recurred across specimens from the same donor (supplementary Figure 1d), finding a median of 6,700 peptides (range: 1,449 -15,264) detected in at least two specimens.

Predicted HLA-I binding affinity varied across specimens (Supplementary Fig. 1e,f). Among colorectal polyps, 15.3% of peptides were predicted binders compared with 55.5% in non-colon tissue (MPNST, Liver metastasis). Because directly observed peptides need not score as predicted binders [25], we retained all dia-PASEF quantified peptides for downstream analysis. Specimens from the same donors had median Jaccard similarity as 0.83, compared with 0.23 across donors, indicating that the peptidome is largely donor-determined. However, the number of peptides identified per donor increased with each additional specimen sampled. This is consistent with incomplete detection at low input rather than with differences between lesions (Supplementary Fig. 1g,h).

We next asked whether HLA-I presentation itself changes during polyposis. In RNA-seq from the same cohort, HLA-I genes and β2M were downregulated in polyps (Figure 1e,f, Supplementary Fig. 1i), consistent with early loss of antigen presentation in polyposis [10]. Additionally, β2M loss was also evident at the immunopeptidome level (log2FC = -1.76, p = 0.063).

To identify candidate polyposis-specific antigens, we compared peptide abundance between polyps and mucosa (Figure 2a). We observed that peptides from proteins BZW2, ARHGAP5, MGARP were presented at higher intensity in polyps (Suppl Fig. 2). BZW2 has been reported as overexpressed in colorectal cancer [26], consistent with our differential analysis. Peptides from β-catenin pathway targets known to be upregulated in polyposis, including AXIN2, OLFM4 and ASCL2, were likewise presented at increased intensity (Suppl Fig. 2e). The specific peptides differ between donors, since presentation depends on HLA-I genotype. Nonetheless, the overall direction of change matched these established expression patterns, indicating that immunopeptidomics recovers known polyposis biology at the level of presented peptides.

**Figure 2.**
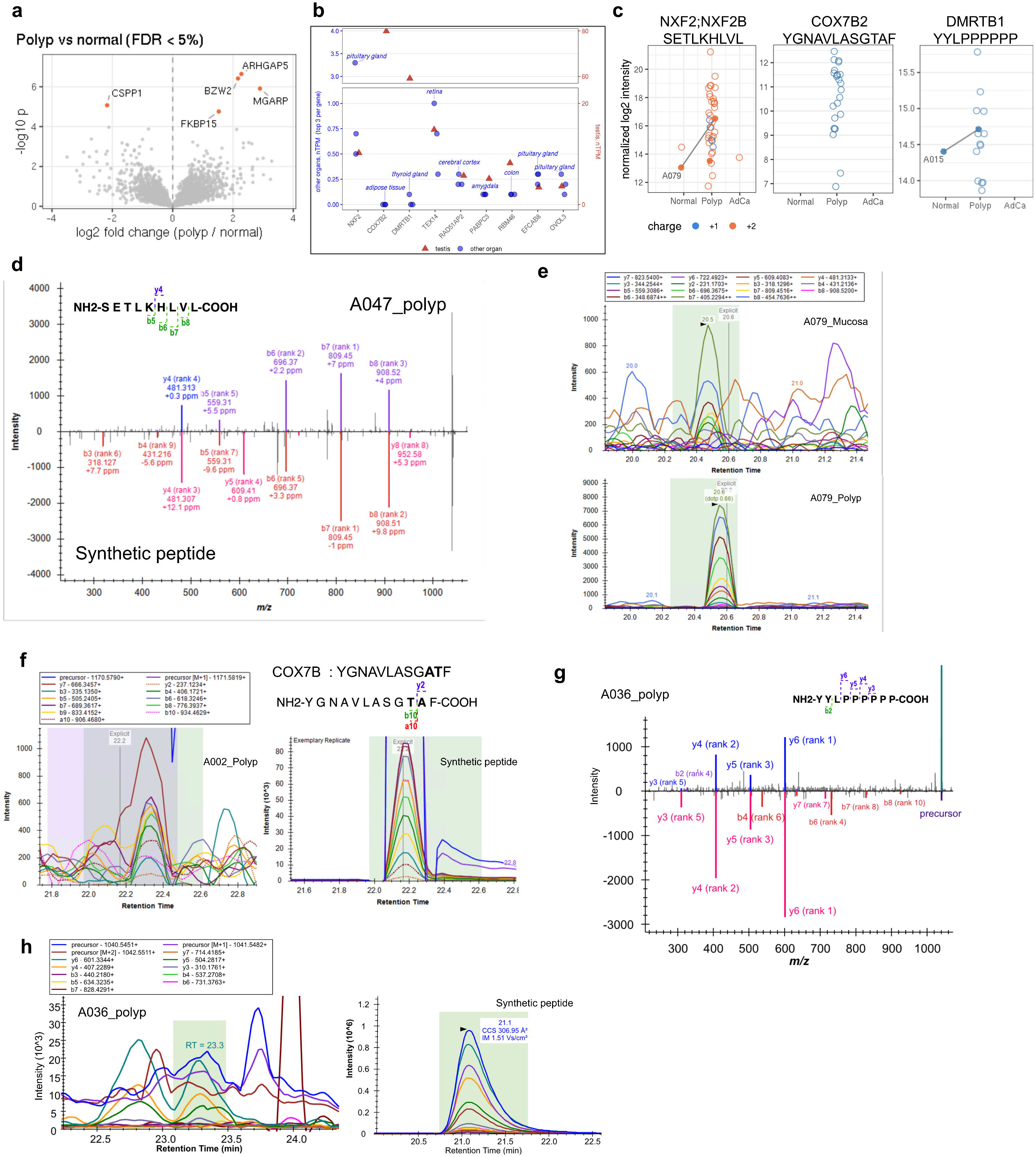
Candidate peptide identification using synthetic standards. **a**) Volcano plot of peptide abundance in polyps versus normal mucosa. Peptides passing FDR < 5% are coloured, with source genes labelled. **b**) Expression of germ-cell genes in normal human tissue (GTEx) in Testis and other top three organs. **c**) Normalised log2 intensity of the three candidate peptides SETLKHLVL (NXF2;NXF2B), YGNAVLASGTAF (COX7B2) and YYLPPPPPP (DMRTB1) across disease stages, coloured by precursor charge state. Specimens with matched mucosa are shown as filled circles. **d**) Mirror plot of the dda-PASEF spectrum for SETLKHLVL(+1) from polyp specimen A047 (Sample ID 97) against the corresponding synthetic peptide standard. Inset shows b- and y-ion assignments on the peptide sequence. **e**) dia-PASEF extracted-ion chromatograms for SETLKHLVL(+2) in matched mucosa and polyp from donor A079 (Sample IDs are 76 and 77). Inset shows the Skyline colour scheme for each fragment ion. **f**) Extracted-ion chromatograms for YGNAVLASGTAF in polyp from donor A002 (Sample ID 20, left) and the synthetic peptide standard (right). Dotted lines mark the b10, y2 and a10 ions that discriminate this COX7B2 peptide from its COX7B paralog product. **g**) Mirror plot of the dda-PASEF spectrum for YYLPPPPPP from polyp of donor A036 (Sample ID 71) against the synthetic peptide standard. **h**) Extracted-ion chromatograms for YYLPPPPPP in polyp (left) and the synthetic peptide standard (right). Inset shows color scheme for precursor ions and its first two isotopes, and for fragment-ions. nTPM: normalized transcripts per million.

### Synthetic peptide standards confirm germ-cell restricted peptide identifications

Given the low tumor mutational burden in polyps, we focused on genes whose expression is normally restricted to immune-privileged organs (Supplementary Table 1). We started with 857 genes (Methods); peptide identification at 1% FDR yielded 349 source proteins. We filtered out peptides that are shared with somatically expressed protein-coding genes, resulting in 107 proteins. Next we kept genes which are immune privileged in both HPA and GTEx (86 proteins). Restricting to peptides of 8-14 amino acids left 58 genes. We then excluded genes expressed above 0.2 TPM in heart, removed isoleucine/leucine-equivalent sequences, and manually selected based on m-score, predicted HLA-I binding and match to synthetic standard spectra, giving nine peptides from germ-cell restricted genes (Table 1, Figure 2b).

**Table 1:** Candidate peptide sequence, source genes and dda-PASEF spectrum charge.

| ID | Sequence | Gene | Uniprot ID | Start | End | Charge |
|---|---|---|---|---|---|---|
| 1 | SETLKHLVL | NXF2;NXF2B | Q9GZY0 | 385 | 393 | +1 |
| 2 | YGNAVLASGTAF | COX7B2 | Q8TF08 | 38 | 49 | +1 |
| 3 | YYLPPPPPP | DMRTB1 | Q96MA1 | 262 | 270 | +1 |
| 4 | EKDISLTDI | TEX14 | Q8IWB6 | 1171 | 1179 | +1 |
| 5 | VYLKEIAKK | RAD51AP2 | Q09MP3 | 273 | 281 | +2 |
| 6 | PAAAAAAAAT | PABPC3 | Q9H361 | 488 | 497 | +1 |
| 7 | SIEVTLAK | RBM46 | Q8TBY0 | 300 | 307 | +1 |
| 8 | MNLVAVASTRQKID | EFCAB8 | A8MWE9 | 212 | 225 | +2 |
| 9 | DAYIPGGPL | OVOL3 | A0AA75L0V4 | 27 | 35 | +1 |

We next examined where the source genes are expressed outside testis, and in immune-accessible tissues. NXF2 is expressed in epididymis (14.3 nTPM) and at lower levels in pituitary glands (3.3 nTPM), and COX7B2 shows minimal placental expression (1.1 nTPM). Epididymis and placenta are also both immune-privileged. All remaining source genes were below 1 nTPM in every other tissue.

Quantification of three candidate peptides is shown in Figure 2c. For each, we confirmed the polyp-derived spectrum against a synthetic peptide standard in water. Figure 2d shows a DDA spectrum of SETLKHLVL/1 from NXF2;NXF2B gene, also known as cancer-testis antigen CT39, identified in polyp against the synthetic peptide, with coverage of both b-and y-ion series. This peptide was presented at approximately eight-fold higher intensity in polyp than in matched mucosa (Figure 2e).

In the figure 2f, we show the DIA MS/MS chromatogram for YGNAVLASG<u>TA</u>F/1 from COX7B2. This gene has a paralog COX7B which is broadly expressed across human tissues and yields an isobaric peptide YGNAVLASG<u>AT</u>F. We therefore used the *b10*, *y2* and *a10* ions, which are specific to the COX7B2 sequence, to confirm that the COX7B2-derived peptide was present. These three ions were detected in both the polyp and synthetic peptide chromatograms. Their summed intensity was used to quantify the COX7B2 peptide (Figure 2c).

Next, we show the spectrum for peptide YYLPPPPPP/1 from DMRTB1 (Figure 2g). Poly-proline peptides characteristically give broad chromatographic peaks, which was also observed in the DIA chromatograms (Figure 2h). The top fragment-ions were from poly-proline regions (y6, y4, y5, y3 ions), and b-ions were comparatively weak, hence we used the MS1 isotope correlation and MS1 isotope overlap scores from OpenSWATH to confirm peak assignment. The sum of MS1-precursor and first isotope was used for quantification (Figure 2c).

The peptide DAYIPGGPL/1 is derived from the OVOL3 protein isoform A0AA75L0V4, which results from retention of a 72-nt intron at chr19:36,111,297 -36,111,368. The comparison against synthetic standards and intensity response for remaining candidates are shown in Supplementary Figure 4 and 5. RAD51AP2 was elevated in normal mucosa of donor A002, which showed reduced B2M in the same tissue (Fig. 1f), suggesting that normal mucosa from colectomy specimens is an imperfect reference for healthy colon. None of the peptides were reported in the HLA Ligand Atlas [8]. Peptides from OVOL3 and EFCAB8 were also detected in the normal autopsy specimen B004; however, their expression was elevated in polyps.

### Source genes are expressed at low levels in colorectal tumours

We next asked whether there is transcript-level evidence for these genes, using colon adenocarcinoma (COAD) and rectal adenocarcinoma (READ) data from TCGA. The NXF2;NXF2B locus was expressed in 34.2% of tumour specimens. Other genes COX7B2 and PABPC3 were also significantly higher in tumours than in normal tissue (Fig. 3a). RAD51AP2, EFCAB8 and OVOL3 followed the same pattern, while DMRTB1, TEX14 and RBM46 trended higher in tumours without reaching significance. Next, in our own FAP RNA-seq, eight of the nine genes were detected in a higher fraction of adenocarcinoma samples than of matched normal tissue (Suppl Fig. 6a), six of which are shown in Figure 3b. NXF2 and DMRTB1 are depicted in Suppl Fig. 6b, and COX7B2 was not detected in this cohort.

**Figure 3.**
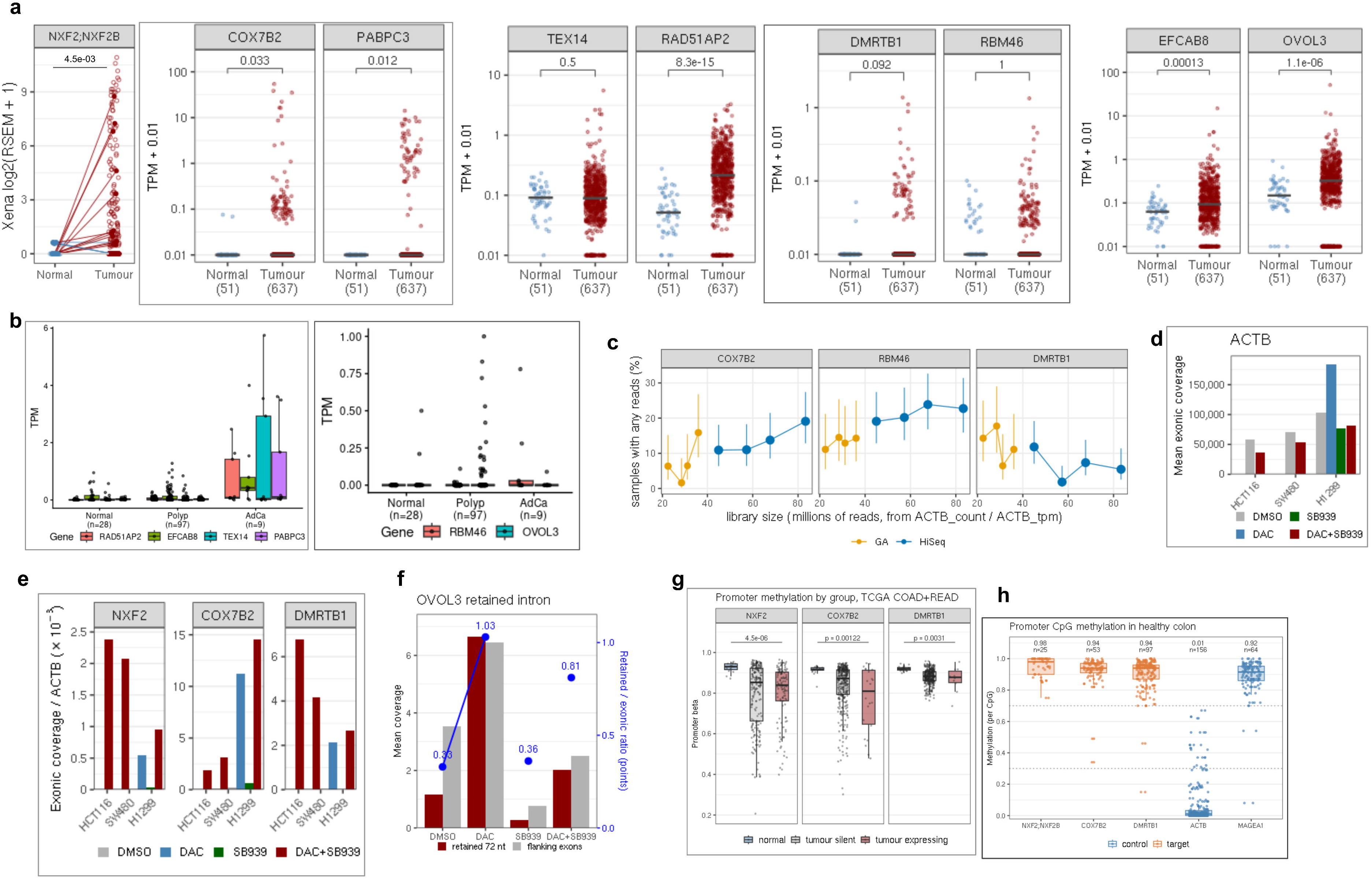
Germ-cell-restricted loci are silenced by DNA methylation in normal colon. **a)** Expression of genes from TCGA COAD and READ. Left, NXF2;NXF2B in matched normal–tumour pairs; lines connect pairs and are coloured by direction of change, with p-values from Wilcoxon signed-rank test. Next, remaining eight genes with GDC STAR TPM, with p-value using Mann-Whitney U test. **b)** STAR TPM in FAP polyps. Left, RAD51AP2, EFCAB8, TEX14, and PABPC3. Right, RBM46 and OVOL3. **c)** Percentage of TCGA samples with any reads for COX7B2, RBM46 and DMRTB1 as a function of library size, estimated from ACTB counts and TPM. Points are binned by library size and split by sequencing platform Illumina Genome analyzer (GA) and HiSeq; error bars show Wilson 95% confidence intervals. **d)** Mean exonic coverage of ACTB in HCT116, SW480 and H1299 cells treated with DMSO, DAC, SB939, or DAC+SB939 (GSE209772), shown as a loading control for (e). **e**) Exonic coverage of NXF2, COX7B2 and DMRTB1 normalised to ACTB. **f**) Coverage of the OVOL3 retained 72-nt intron and its flanking exons in H1299 cells (bars, left axis), with the retained-to-exonic ratio overlaid (points, right axis). **g**) Promoter CpG methylation (Illumina 450K β values) for NXF2, COX7B2 and DMRTB1 in TCGA COAD and READ, split into normal tissue, tumours not expressing the gene, and tumours expressing the gene. p-values from unpaired Wilcoxon rank-sum test. **h**) Per-CpG promoter methylation in healthy colon (Roadmap E106 WGBS) for the three target loci and for the controls ACTB and MAGEA1. Numbers above each group give the median methylation and the number of CpGs assessed. DAC: Decitabine; SB939: histone deacetylase inhibitor pracinostat.

Expression sat below 1 TPM for most of these genes in TCGA, which led us to ask whether detection depends on sequencing depth. Detection of COX7B2 and RBM46 genes correlated positively with depth (Fig. 3c). DMRTB1 showed no such association. It has 60.5% GC content across the 1910-nt transcript (ENST00000371445.3), with a block in exon 1 peaking at 86%. We speculate this could have led to lower detection in bulk-RNAseq dataset. Nonetheless, the normal samples were sequenced more deeply than the tumours (median 75.6 M vs 47.8 M reads, Wilcoxon p = 2.3e-06 post instrument-stratification; library size taken as ACTB count / ACTB TPM).

We were surprised that peptides from source genes expressed at such low TPM could be presented and detected with LC-MS/MS. Other studies using targeted PRM acquisition have detected HLA-I peptides from transcripts expressed as low as 0.02 FPKM (Supplementary Table 2) [27]. The MAGE-A4-directed TCR used in the approved synovial sarcoma therapy also cross-reacts with MAGE-A8 peptide [28], which had an expression of 0.2 TPM [29]. Low transcript abundance is therefore compatible with surface presentation and, importantly, with direct detection by highly sensitive HLA-peptide enrichment and high sensitivity LC-MS/MS workflows. This highlights the ability of modern immunopeptidomics that cannot be reliably inferred from transcript abundance alone.

### Loss of DNA methylation derepresses these loci in colorectal cancer

We next asked what drives transcription of these germ-cell enriched genes. Several cancer testis genes are epigenetically silenced in somatic tissue [11], and global hypomethylation is a recognised feature of colorectal cancer [23,30]. We therefore examined their expression in cell lines treated (GSE209772) with the DNA methyltransferase inhibitor decitabine (DAC) or the histone deacetylase inhibitor SB939 [31]. Expression of all three genes was induced by DAC, whereas SB939 had virtually no effect (Figure 3d,e). We observed a similar pattern for other source genes TEX14, RBM46, and EFCAB8 (Supplementary Figure 6c). The OVOL3 peptide derives from a retained intron, and in H1299 cells DAC increased coverage of this intron 5.7-fold while coverage of the flanking exons rose only 1.8-fold, raising the retained-to-exonic ratio from 0.33 to 1.03 (Fig. 3f). Surprisingly, OVOL3 was silent in the colorectal lines HCT116 and SW480. The other gene RAD51AP2 did not show much effect in the cell lines (Supplementary Figure 6d).

We next asked whether the same relationship between hypomethylation and expression holds in TCGA colorectal cancer. We compared Illumina 450k β values between normal and tumor samples (probes listed in Supplementary Table 3). Promoter CpG methylation was high in normal colon and significantly lower in tumours for all three genes, and among tumours, those expressing the gene were less methylated than those that were not (Figure 3g). Hypomethylation was also seen at probes for RBM46, EFCAB8, TEX14 (Supplementary Fig. 6e). For PABPC3 and RAD51AP2, individual probes showed significant hypo- and hypermethylation within the same gene, whereas OVOL3 showed hypermethylation at two probes (Supplementary Figure 6f).

Next we also confirmed that these regions are methylated in a non-cancer donor, whole-genome bisulfite sequencing (WGBS) of healthy autopsy sigmoid colon from the Roadmap Epigenomics dataset (E106, composite of two male donors) [32]. WGBS provides denser CpG coverage than the Illumina array, including sites not represented by the 450K probes. All three loci were methylated, as were the remaining source genes with the exception of RAD51AP2 (Fig. 3h, Supplementary Fig. 6g). Studies have demonstrated that its expression is restricted primarily to meiotic tissues, particularly adult testis and fetal ovary [33].

### Loci lose methylation in premalignant lesions

We next examined CpG methylation in polyps using enzymatic methyl-seq (EM-seq), which provides better coverage than WGBS [23]. Methylation at these loci was significantly (p < 0.05) reduced in polyps compared with matched mucosa (Figure 4a). Although this is attributed to global DNA hypomethylation, not all genes follow the same pattern [23]. The COX7B, paralog of COX7B2, showed no-change or slightly higher methylation in polyp (Supplementary Figure 7a). Five of the remaining seven genes showed reduced CpG methylation; RBM46 and OVOL3 did not. RBM46 promoter was nonetheless hypomethylated in TCGA tumour samples (Supplementary Figure 7b,c).

**Figure 4.**
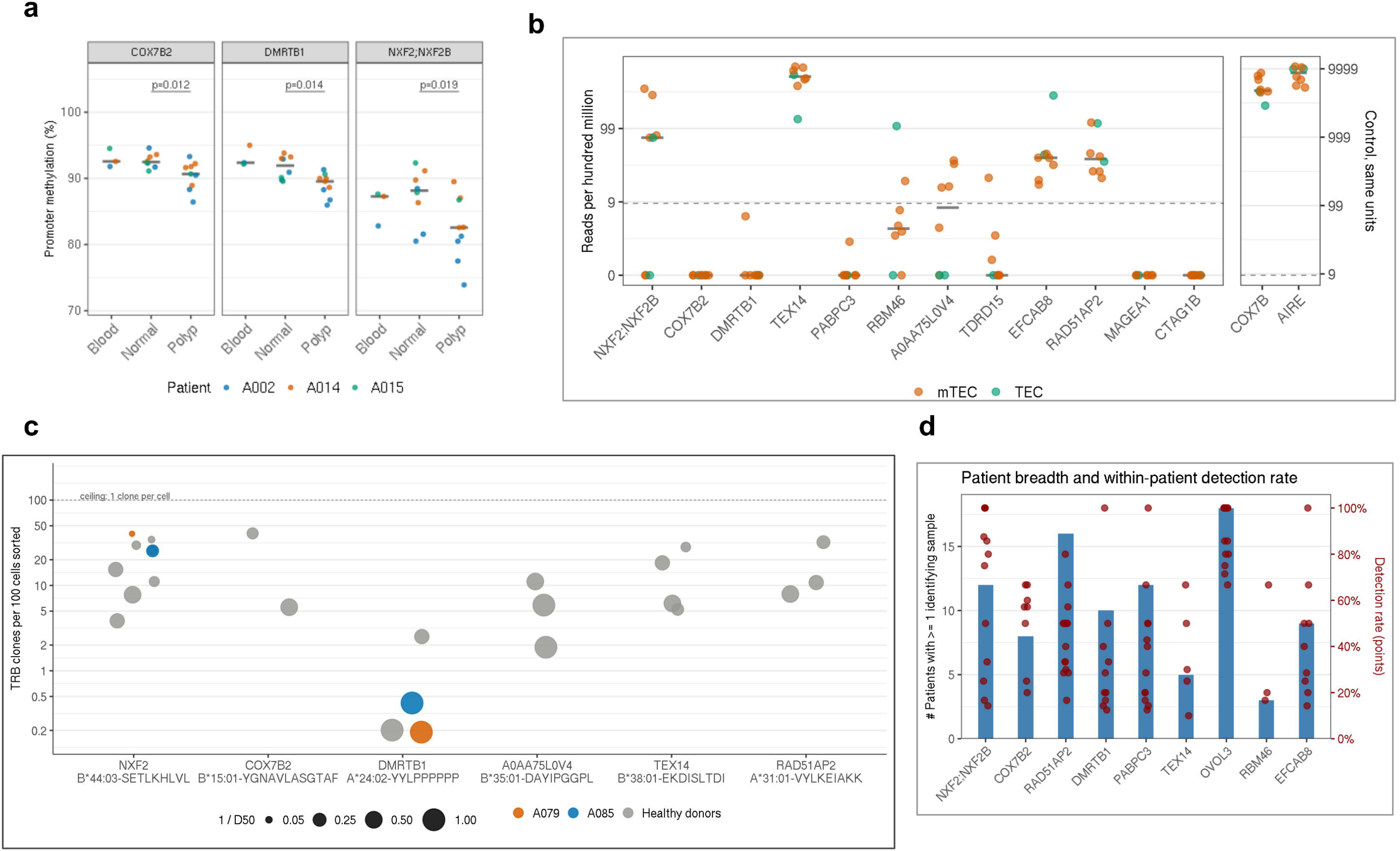
Methylation loss in polyps and T cell recognition of candidate peptides. **a**) Promoter CpG methylation for COX7B2, DMRTB1 and NXF2;NXF2B loci in blood, matched mucosa and polyp from three FAP donors. Each point is one specimen; horizontal bars show the group median. p-values from two-sided within-donor permutation test. **b**) Expression of the ten source genes in medullary thymic epithelial cells (mTECs) and thymic epithelial cells (TECs). Each point is one donor; horizontal bars show the median. Dashed line marks the 8.55 rphm detection threshold (Laumont 2018). MAGEA1 and CTAG1B (NY-ESO-1) are shown as thymically silent cancer-testis antigens; AIRE is shown as a positive control on the right axis. **c**) TCRβ repertoire of peptide-dextramer-sorted T cells. Each point is one donor; the y-axis gives the number of TRB clonotypes recovered per 100 cells sorted, and point size gives 1/D50, where D50 is the fraction of clonotypes accounting for 50% of reads. Larger points indicate more clonally focused repertoires. Dashed line marks the ceiling of one clonotype per cell. Peptides are labelled with their source gene and HLA-I allele of dextramer used. FAP donors are coloured; healthy donors in grey. **d)** Left y-axis, Number of patients with peptide recovered. Right y-axis, Fraction of samples within the same donor the peptide was recovered. rphm: reads per hundred million.

### Thymic expression and circulating repertoires against candidate peptides

A determinant of whether a presented peptide can elicit a T cell response is whether its source gene is expressed in the thymus, where developing T cells undergo negative selection against self-peptides. We therefore examined expression of these genes in medullary thymic epithelial cells (mTECs), the cell type responsible for negative selection [21,34]. NXF2;NXF2B, TEX14, EFCAB8 and RAD51AP2 were expressed above the detection threshold of 8.55 rphm (Fig. 4b), indicating that T cells recognising their peptides are likely subject to central tolerance. In contrast, COX7B2, DMRTB1 and PABPC3 were not detected, nor were the known cancer testis antigens MAGE-A1 and NY-ESO-1 (CTAG1B), suggesting that peptides from these genes may escape central tolerance. RBM46 and OVOL3 (A0AA75L0V4) were detected near the threshold, with expression varying between donors.

To test whether these peptides are recognised by T cells, we sorted peptide-dextramer-binding T cells (Supplementary Table) and performed bulk TCRβ sequencing. Dextramer-binding cells differed markedly in repertoire structure (Fig. 4c). For NXF2, TEX14, RAD51AP2 sorted cells comprised many distinct clones at low individual frequency, consistent with low-avidity or background binding. Although we observed the NXF2 peptide about eight-fold high in polyps of A079 (Figure 2e), we did not observe any clonal expansion in PBMCs of A079.

DMRTB1 behaved differently. Only a handful of clones dominated the sort in both FAP patients and one healthy donor, indicating clonal dominance rather than polyclonal binding. The A0AA75L0V4 peptide showed the same pattern, though less pronounced. Despite COX7B2 being undetected in mTECs, we did not recover a comparable signal. The COX7B2 peptide differs from its COX7B counterpart only at positions 10 and 11, where threonine and alanine are transposed (G<u>TA</u>F versus G<u>AT</u>F), so high-avidity clones recognising COX7B2 may already have been selected against the broadly expressed paralog.

No TCRβ motif was shared across donors for these peptides. Shared, or public, motifs are typical of viral epitopes such as influenza but are rarely found for self-antigens (Supplementary Fig. 7d). Repertoire structure rather than motif sharing is therefore the more informative readout of recognition here.

## Discussion

We identify HLA-I peptides from germ-cell-restricted genes presented on premalignant colorectal lesions. Seven of these peptides are absent from healthy colon in both the HLA Ligand Atlas and autopsy specimens investigated in this study, and their presentation tracks with demethylation at loci and transcript detection across cell lines, tumours and polyps. It is known that at the start of polyposis the HLA-I presentation pathway declines, NK cells that survey HLA-I are reduced, and exhausted T cells accumulate by the adenocarcinoma stage [35]. Alterations in the presented peptidome, however, were unknown. We show that germ-cell specific peptides are presented at this stage, so the antigenic change accompanies the immune changes already described.

Therapies directed at peptide–HLA class I complexes have entered clinical practice. Tebentafusp engages T cells against an HLA-A*02:01–restricted gp100 peptide in uveal melanoma [36], and engineered T cells specific for an HLA-A*02:01–restricted MAGE-A4 peptide have been approved for synovial sarcoma [37]. Both show that a single presented peptide can be a tractable target in solid tumours. Notably, gp100 (PMEL) and MART-1 (MLANA) are melanocyte differentiation antigens expressed in normal tissue, so detection in normal tissue does not by itself disqualify a candidate. On this basis the peptides we detected from OVOL3 and EFCAB8 in a normal autopsy specimen remain of interest, particularly as both derive from germ-cell-restricted genes. OVOL3 showed no methylation change at its promoter in either the polyp or TCGA data. Decitabine nonetheless increased its intron-retained transcript ENST00000585332 (Fig. 3d), and the peptide derives from that transcript, suggesting that the effect is not mediated by promoter methylation. Whether it reflects demethylation elsewhere in the locus, or regulation by other factors, remains unclear.

Because β2M is already reduced in polyps and falls further in carcinomas, peptides presented at the polyp stage may be largely absent by the time a tumour is resected, which could explain why they were not found in established CRC datasets. Timing is already recognised as a determinant of anti-PD-L1 response, which depends on when treatment is given relative to the immunosuppressive state of the tumour [38]. The precancerous window may be the more tractable one for the same reason. Although carcinomas carry high mutational and neoantigen burdens, they converge on an immunosuppressive environment that would blunt any intervention [39].

Transcript-level evidence for these genes was sparse, and detection scaled with library size for several of them. Targeted sequencing would be required to measure these transcripts reliably across stages. DMRTB1 was the exception, showing no association with depth; its high GC content, with a block in exon 1 reaching 86%, offers a plausible explanation for its poor recovery. More generally, expression-based screening is not a substitute for measuring the peptidome directly.

Seven of the nine peptides were identified from +1 charge state spectra, although quantification used both +1 and +2 charge states (Table 1). These identifications are possible because PASEF both removes interfering ions from each scan and increases sensitivity by condensing the ion signal [40].

Candidate peptides were confirmed against synthetic standards. Retention times differed between polyp and synthetic chromatograms because the standards were prepared in water rather than in tissue matrix; fragment ion identity is therefore the basis of confirmation. We also detected low-level signals for some germ-cell restricted peptides in matched mucosa, where we had expected none. This most likely reflects a field effect, since mucosa was collected at colectomy from patients with extensive polyposis and some inflammation is expected. The HLA Ligand Atlas, which covers benign tissue from 16 donors, is therefore the more appropriate negative reference, and these peptides were absent from it.

Candidate antigens are conventionally prioritised by predicted HLA-I binding affinity, on the reasoning that mutations strengthen peptide loading for neoantigens. That reasoning holds for neoantigens [1, 41] as well as self-antigens [42], but our data suggest that besides the binding prediction, thymic expression of the source gene is an equally useful criterion. We condensed these results into a convergence table (Supplementary Table 9). The COX7B2 result illustrates this principle, as the peptide differs from its ubiquitously expressed paralog COX7B by a transposition of two residues, and we did not recover as strong a clonal repertoire as we did for DMRTB1 (Figure 4b,c). Peptide prioritization matters practically, since cognate T cells may be present at frequencies near 0.1% and dextramer assays require on the order of 10 cells per peptide, so only a handful of candidates can be tested from a standard blood draw.

Although sensitive immuno-peptidomics allows the identification of interesting antigens from small polyp samples, their coverage still needs to be addressed (Figure 4d). Specimens were not homogenized, so sampling depth and between-lesion heterogeneity cannot be separated. Additionally, cognate clones were identified in blood, but circulating and tissue-resident repertoires overlap only partially, so we cannot say whether these T cells reach the lesion. Whether these antigens appear in sporadic adenomas, and whether cognate T cells reach the lesion itself, can now be tested directly.

Our study places the emergence of an altered antigenic landscape among the early molecular events of colorectal tumorigenesis. Germ-cell-restricted loci that are normally silenced are hypomethylated in premalignant lesions, and their products reach the cell surface at transcript abundances that would not have nominated them as candidates. Whether the same antigens appear in sporadic lesions, and whether tissue-resident T cells encounter them, can now be addressed directly.

## Acknowledgement

This project is one of the Human Tumor Atlas Network (HTAN) Projects and was supported by NCI grant no. U2CCA233311 to MPS and JMF. SG is grateful for the support of the Stanford Center for Computational, Evolutionary, and Human Genomics through the CEHG Postdoctoral Fellowship. SK and OKK were supported by European Union EIC Transition grant no. 101112948 (PeptiCHIP).

## Competing interests

MPS is a cofounder and scientific advisor of Crosshair Therapeutics, Exposomics, Filtricine, Fodsel, Iollo, InVu Health, January AI, Marble Therapeutics, Mirvie, Next Thought AI, Orange Street Ventures, Personalis, Protos Biologics, Qbio, RTHM, and SensOmics, and is a scientific advisor to Abbratech, Applied Cognition, Enovone, Jupiter Therapeutics, M3 Helium, Mitrix, Neuvivo, Onza, Sigil Biosciences, TranscribeGlass, WndrHLTH, and Yuvan Research. RS and DH are employees of Bruker Daltonics. SK and OKK were employees of ValoTx. SG and MPS are inventors on a provisional patent application covering the use of the peptides described in this manuscript.

## List of Supplementary Tables

Supplementary Table 1. Tumour mutational burden from FAP specimens.

Supplementary Table 2. HLA-I peptide and gene FPKM.

Supplementary Table 3. Illumina HumanMethylation450 probes used for promoter methylation analysis.

Supplementary Table 5. 10x Genomics Single-Cell TCR Repertoire From Five Antigens (EBV, Influenza, MAGE-A1, NY-ESO-1).

Supplementary Table 7. Cohort donor and their HLA-I alleles.

Supplementary Table 8. HLA class I genotypes of Stanford Blood Center donors.

Supplementary Table 9. Summary of evidence for the nine candidate peptides across independent assays and cohorts.

Supplementary Table 10. Cohort summary.

Supplementary Table 11. Sample Summary.

## List of Supplementary Figures

Supplementary Figure 1. Cohort composition, acquisition-mode comparison, and depth of the immunopeptidome dataset.

Supplementary Figure 2. dia-PASEF quantitation for all peptides of selected source genes.

Supplementary Figure 3. Detection of candidate peptides across donors and specimens.

Supplementary Figure 4. Spectral and chromatographic evidence for candidate peptides against synthetic standards.

Supplementary Figure 5. Normalised log2 intensity of remaining six candidate peptides across disease stages.

Supplementary Figure 6. Transcript detection, epigenetic reactivation and promoter methylation of the remaining source genes.

Supplementary Figure 7. Promoter methylation in polyps and reference T cell and thymic data.

## Methods

### Specimen collection

Colorectal polyp specimens were collected from patients undergoing colonoscopy procedures at Stanford Hospital under IRB protocol #47044. For a subset of patients undergoing colectomy, matched normal colorectal tissue specimens were also collected. Following excision, specimen dimensions were recorded, and tissue designated for immunopeptidomics analysis was fresh-frozen in liquid nitrogen and stored until mass spectrometry processing. Sorted cells were collected directly into the lysis buffer and stored at −80 °C until further processing.

### HLA-I peptide isolation

We collected HLA-I immunopeptidomes by lysing tissue, isolating HLA-I–bound peptides on PeptiCHIP cartridges coated with the pan-HLA-I antibody W6/32, and acquiring mass spectra on a high sensitivity Bruker timsTOF Ultra 2 [24].

Tissue samples were lysed in 200 µL of ice-cold lysis buffer containing 150 mM NaCl, 50 mM Tris-HCl pH 7.4, 1% IGEPAL CA-630, and one Pierce protease inhibitor tablet. The lysate was slowly mixed at 4°C for 1 hour(s) with a vertical rotation mixer. A 10 µL aliquot was reserved for proteomics analysis. The remaining lysate was centrifuged at 500 × g for 10 minutes at 4°C. The supernatant was transferred to a new tube and further centrifuged at 16,000 × g for 30 minutes at 4°C.

The supernatant was concentrated using a 3k centrifugal MWCO filter by spinning at 14,000 x g for 30 mins 4°C. The concentrate was collected and used for HLA-I complex capture using the pan-HLA-I antibody W6/32 on the PeptiCHIP platform. HLA-I-peptide complexes were immunoaffinity-purified using PeptiCHIP microfluidic thiolene chips. These microchips, featuring an array of ∼14,400 micropillars to increase surface area, were pre-coated with biotin-PEG 4-alkyne (Sigma-Aldrich, St. Louis, MO). The final chip functionalization was performed immediately before loading the samples by pre-coating the biotin-functionalized microchips with streptavidin (0.1 mg/mL in 1× PBS, Sigma-Aldrich, St. Louis, MO, USA), preconditioning with BSA (100 μg/mL in 15 mM PBS, Sigma-Aldrich, St. Louis, MO), and immobilizing the biotinylated anti-panHLA antibody (clone W6/32, BioLegend, San Diego, CA, USA). HLA-I complexes were captured by adding the lysates to the antibody-coupled PeptiCHIPs. The peptide-HLA-complexes were eluted from PeptiCHIPs with 7% acetic acid (Fisher Chemical, Waltham, MA, USA) in 50% methanol (Fisher Chemical, Waltham, MA, USA).

The eluted samples were then desalted using Sep-Pak tC18 cartridges (Waters, Milford, MA, USA). The cartridges were prewashed with 80% acetonitrile (ACN) in 0.1% trifluoroacetic acid (TFA), followed by washing with 0.1% TFA. Peptides were purified from the HLA-I complexes by loading them into cartridges and eluting with 30% ACN in 0.1% TFA. The samples were further purified using 10K Amicon Ultra spin filters (Merck Millipore, Billerica, MA, USA) and were finally dried using vacuum centrifugation. The isolated peptides were stored at -80°C until mass spectrometry acquisition.

### Mass spectrometry data acquisition

Peptides were analyzed by liquid chromatography-mass spectrometry (LC-MS), with the following conditions: Nanoflow reversed-phase liquid chromatography was performed on a nanoElute2 system (Bruker Daltonics) connected to a timsTOF Ultra2 mass spectrometer (Bruker Daltonics) via a CaptiveSpray nano-electrospray source. Peptides were separated using a 30 min gradient on a C18 column (25cm × 75 μm, 1.7 μm, IonOpticks). The mobile phase A was water with 0.1 vol% formic acid and mobile phase B was ACN with 0.1 vol% formic acid. Peptides were separated with a linear and stepped gradient ranging from 5 to 23% solvent B over 18 min, 23 to 35% in 4 min, 35 to 90% in 4 min, and held at 90% for 4 min at 250 nl/min. The mass spectrometer was operated in ddaPASEF and diaPASEF mode. For ddaPASEF analysis, MS scans were acquired from 100 to 1700 m/z and 1/K0 = 0.7 Vs cm−2 to 1.7 Vs cm−2. Seven PASEF ramps were acquired with an accumulation and ramp time of 200 ms and total cycle time of 1.66s. Precursors above the minimum intensity threshold of 1000 were isolated with 2 Th at < 700 m/z or 3 Th >800 m/z and reacquired until a target intensity of 14,000. Active exclusion was set to 24s, and the collision energy (CE) was a linear function between the CE and the ion-mobility starting from 15 eV at 1/K0 = 0.6 Vs cm−2 to 57 eV at 1/K0 = 1.6 V cm−2. For diaPASEF analysis, an optimized isolation window scheme in the m/z versus ion mobility plane was designed using pydiAID that comprised MS scans from 100 to 1700 m/z and 1/K0 = 0.69 Vs cm−2 to 1.70 Vs cm−2. that covered the range of 347 to 1265 m/z and an ion mobility range of 0.69 -1.70 Vs cm−2, with variable isolation windows (no overlap) with a total cycle time of 1.66 s (Table1). The collision energy (CE) was a linear function between the CE and the ion-mobility starting from 19.20 eV at 1/K0 = 0.70 Vs cm−2 to 34.32 eV at 1/K0 = 1.06 Vs cm−2 to 36.00 eV at 1/K0 = 1.10 Vs cm−2 to 46.08 eV at 1/K0 = 1.34 Vs cm−2 to 60.36 eV at 1/K0 = 1.68 V cm−2. Accumulation and ramp time were set to 121 ms. The instrument was operated in high sensitivity mode for both acquisitions. The acquisition scheme for DIA was optimized using <u>pydiaid</u> [43]. The window scheme is in Supplementary Table.

### Mass spectrometry data analysis

The DDA data was analyzed using FragPipe (v22.0) with MSFragger (v4.1) for peptide identification [44]. The reference proteome from UniProtKB/Swiss-Prot for Homo sapiens (TaxonID 9606; downloaded 19 October 2024) was searched using “Non-specific HLA” workflow with a precursor mass tolerance of 15 ppm and a fragment mass tolerance of 15 ppm. Variable modifications included methionine oxidation (+15.995 Da), protein N-terminal acetylation (+42.011 Da), N-terminal pyroglutamate formation from glutamine (−17.027 Da), and N-terminal pyroglutamate formation from glutamic acid (−18.011 Da) with a maximum of 3 variable modifications per peptide. Charge states were restricted to 1–4 and peptide length filters were set to 7–15 amino acids. For database search, the MSBooster was enabled with RT model set to DIA-NN, and spectra model as “Prosit_2023_intensity_timsTOF” [45]. The ion, PSM and peptide FDR filter were set to 10%. Spectral libraries were generated for each sample using SpeclibGen.

The DIA data was analyzed using OpenSWATH workflow (OpenMS v3.4.1) [46]. For this, the spectral libraries were merged from all runs and decoys were added using OpenSwathDecoyGenerator with -switchKR flag set to false. The Bruker .d files were converted to mzML using MSConvert with -- combineIonMobilitySpectra flag [47]. Each DIA file was processed using OpenSWATH, followed by statistical scoring with PyProphet (v3.0.0). Peptide identifications were filtered at per-patient level with 1% global FDR [48].

Candidate peptides were manually reviewed in Skyline [49]. For NXF2, COX7B2, PABPC3 and RAD51AP2, spectral library coordinates were imported as a transition list, chromatographic peak boundaries were manually curated, and non-quantitative fragment ions were excluded from downstream quantification.

*Immune-privileged gene panel*:

Candidate source genes were assembled from Human Protein Atlas consensus RNA data, using normalised transcripts per million (nTPM) [50]. The immune-privileged compartment was defined as testis, placenta, epididymis and seminal vesicle. Genes were admitted through three layers, and the panel is their union. Layer A required ≥3 nTPM in the privileged compartment together with <1 nTPM in every somatic tissue (536 genes). Layer B relaxed the somatic ceiling to <5 nTPM while requiring <1 nTPM in all vital organs (578 genes). Layer C added a curated CTdatabase from http://www.cta.lncc.br. The layers overlap substantially; the union comprises 857 distinct genes.

### HLA Typing

HLA-I allotypes were determined by applying OptiType and arcasHLA to WGS, bulk-RNAseq and single-cell RNAseq data [51]. The alleles are given in Supplementary Table 7.

HLA-I peptide binding prediction:

Binding was predicted with NetMHCpan v4.1 [52]. For each peptide–allele pair we took the final rank as the minimum of the binding affinity (BA) and eluted ligand (EL) percentile ranks.

### Cancer Testis Antigen (CTA) identification

Per-tissue expression for the gene panel was taken from Human Protein Atlas consensus files derived from GTEx (rna_tissue_gtex.tsv, 36 tissues), which report nTPM (cross-sample normalised) per gene per tissue. List of CTAs were downloaded from the <u>CTdatabase</u>. Identified CTA source genes detected in polyps were further evaluated for RNA expression across normal tissues using the Human Protein Atlas [50] and <u>GTEx datasets</u> [17]. The corresponding peptide sequences were also queried against the <u>HLA ligand Atlas</u> to assess whether they had been previously detected among naturally presented HLA ligands in benign human tissues [8]. GTEx and HLA ligand Atlas were acquired from autopsy samples, hence represent data from cancer-free participants.

### Peptide reactive T cell isolation using dextramer

HLA-I dextramers were obtained from Immudex and loaded with the peptides according to the manufacturer’s protocol. PBMCs were isolated from donor blood by Ficoll and, where specified, cultured with the peptide of interest for 10 days. Cells were plated in 24-well plates at 1.5M cells/mL in RPMI supplemented with 10% human AB serum with 10 uM peptide. On Day 3, IL-2 (25 U/mL), IL-7 (5 ng/mL) and IL-15 (5 ng/mL) were added. The media was replaced after every 2 days with fresh cytokines. CD8+ T cells were then isolated by magnetic negative enrichment (EasySep, Catalog# 19051). 500k CD8+ cells were incubated with PE-conjugated dextramer following the manufacturer’s protocol, stained with DAPI, and bulk-sorted on a FACSAria Fusion as PE+ DAPI-cells into 96-well plates containing lysis buffer (Takara, Catalog# 635013). The HLA alleles of blood donors are in Supplementary Table 8.

#### bulk-TCR sequencing on dextramer-sorted cells

Full length cDNA was synthesized using SMART-seq chemistry, and TCR sequences were recovered by Han’s nested PCR method [14]. Cells were sorted directly to the lysis buffer (refer to the Takara recipe). First-strand cDNA synthesis directly from cells was primed by the 3’ SMART-Seq CDS Primer II A and uses the SMART-Seq HT Oligonucleotide for template switching at the 5’ end of the transcript. Full length, amplified cDNA was synthesized using a SMART-Seq mRNA HT kit (Cat. #634796). TCR sequence was obtained by a series of two nested PCR reactions. A 4-μl aliquot of the amplified cDNA was used as a template for a second 15 -μl PCR using HotStarTaq DNA polymerase (Qiagen) for TCR sequencing. A 1-μl aliquot of the second PCR product was used as a template for the third 11-μl PCR reaction, which incorporated barcodes and enabled sequencing on the Illumina MiSeq platform and Element AVITI sequencer. The fastq files were used to run a customized TCRseq informatics pipeline, which assigned reads to wells, assigned TCR Va, Ja, Vb, and Jb usage, along with alternate Va/Ja where applicable, and reported CDR3a and CDR3b sequences. This assay was performed by the Human Immune Monitoring Center at Stanford University. This method was based by Han et al [53].

### TCR analysis

Clonotypes were sorted by abundance and a minor sequence merged into a more abundant one if it differed at exactly one amino acid position and its count was below 5% of the parent, with merged reads added to the parent. Plate-wells were aggregated into sorts by summing read counts across wells and barcode plates for each combination (sort, chain, CDR3, V, J). Clonotypes were retained if they had at least 6 reads per well, at least 500 pooled reads, a within-sort frequency of at least 0.002, and were present on at least one plate.

Clonotypes shared between sorts were adjudicated pair by pair. Each shared pair was classified using (i) the frequency ratio between the two sorts, (ii) the dispersion of that ratio across all clones shared by the pair, (iii) plate geometry and sort date, and (iv) whether the two sorts shared an antigen and a restricting allele. A tight ratio across many clones (log SD 0.26-0.36) indicates a single bulk transfer and identifies a clear source; a ratio near 1 with high dispersion (log SD >1.5) indicates two sorts that are not independent, with no resolvable direction. Clones were not removed where the two sorts shared both antigen and dextramer allele, since a public or cross-reactive receptor is then possible. The curated table contains 1,202 TRB and 1,242 TRA clonotypes across 38 sorts.

### Thymic epithelial cell RNA-seq

Raw reads for six sorted mTEC and two unfractionated TEC samples (GSE127825, GSE127826) were downloaded [21,34]. Reads were aligned with STAR 2.7.3a to the GRCh38 primary assembly with GENCODE v47, sjdbOverhang 75, using --quantMode GeneCounts and otherwise default filters. All eight samples were resolved as reverse-stranded. The NXF2;NXF2B locus was quantified with salmon and summed over both copies. The intron-retained OVOL3 isoform A0AA75L0V4 was quantified using ENST00000585332, which retains the 72 nt at chr19:36,111,297– 36,111,368 in frame.

### Roadmap whole-genome bisulfite data

Normal colon methylation was taken from Roadmap Epigenomics fractional-methylation <u>bigWigs</u>; E106 was the only colon sample with WGBS coverage. Tracks are hg19 [32]. Promoter and gene-body windows were derived from TxDb.Hsapiens.UCSC.hg19.knownGene and cross-checked against hg38 windows lifted over with rtracklayer::liftOver; lifted regions that fragmented or split across chromosomes were rejected in favour of native hg19 coordinates. Coverage was summarised as the fraction of reference CpGs covered. the denominator being all CpGs in the window from BSgenome.Hsapiens.UCSC.hg19 rather than only those observed, so that sparse coverage cannot be mistaken for confident methylation. Values plotted are per-CpG fractional methylation. The bedGraph stores two records per CpG, one per strand, so CpG counts are the record count halved.

#### TCGA RNA-seq

Expression data for colon (COAD) and rectal (READ) adenocarcinoma were obtained from the UCSC Xena platform. Gene-level STAR counts and TPM were retrieved from the GDC Xena hub (https://gdc.xenahubs.net). Reads spanning each gene’s own annotated introns were queried from Snaptron (https://snaptron.cs.jhu.edu/tcgav2, recount3 TCGA, STAR, hg38). Junction counts were additionally normalised per 1,000 ACTB junction reads. NXF2;NXF2B locus is quantified only by Xena RSEM. PABPC3, which shares 2.6% of its 76-mers with the far more abundant PABPC1, is reported as excess over estimated leakage: TPM minus 5 × median(PABPC3/PABPC1) × PABPC1 TPM, floored at zero.

### TCGA paired methylation and expression

Illumina Infinium HumanMethylation450 β values were retrieved from the Xena hub. Promoter probes were selected per gene using the IlluminaHumanMethylation450kanno.ilmn12.hg19 annotation, retaining probes annotated TSS1500, TSS200, 1^st^ Exon or 5’ UTR for the target gene; a locus-level β was computed as the mean over that gene’s probes. Methylation and expression were joined on the 12-character TCGA patient barcode after stripping the sample-type suffix.

### Cell line RNA-seq and epigenetic reactivation

Reactivation of the gene panel by DNA methyltransferase and histone deacetylase inhibition was assessed in GSE209772 [31]. Coverage bigWigs were downloaded from GEO with GEOquery::getGEOSuppFiles(), one file per cell line × treatment arm. All GSE209772 bigWigs are hg19, so gene models were taken from TxDb.Hsapiens.UCSC.hg19.knownGene. Per-gene signal was computed as mean exonic coverage (rtracklayer::import.bw restricted to the gene’s exonic intervals), then expressed relative to ACTB mean exonic coverage in the same file, giving a within-sample normalised value.

## Notes

https://www.ncbi.nlm.nih.gov/geo/query/acc.cgi?acc=GSE127825

https://egg2.wustl.edu/roadmap/data/byDataType/dnamethylation/WGBS/FractionalMethylation_bigwig/

https://www.proteinatlas.org/about/download

https://tcga.xenahubs.net

https://gdc.xenahubs.net

https://www.ncbi.nlm.nih.gov/geo/query/acc.cgi?acc=GSE209772

### Summary of Updates

Updated corresponding author list; added figures.

