## Supplementary_Note_Figures for "Germ-cell restricted antigens are presented on HLA-I in premalignant colorectal lesions"

Corresponding authors:

**Supplementary Note 1. Prior evidence for candidate antigens from IEDB**

To assess whether the candidate antigens had been reported previously, we queried the MHC ligand database of the Immune Epitope Database (Vita 2024).

1. NXF2/NXF2B — SETLKHLVL. This peptide was reported in six publications. Five identified it in lymphoid or patient-derived cancer cell lines, which do not represent healthy normal tissue. The sixth reported it in adjacent normal colorectal biopsy tissue; however, the donor had metastatic disease (Neidert 2018), so this tissue cannot be considered a healthy normal reference in the sense of the HLA Ligand Atlas. Together these results indicate that peptide is pre-cancerous tissue specific and a good target for therapeutics. The peptide is shared by NXF5, which is however is considered as a pseudogene.
2. COX7B2 — YGNAVLASGTAF. Not reported in IEDB. However, the corresponding peptide from its paralog COX7B, YGNAVLASGATF, has been observed. The two peptides differ at two positions and share closely related flanking sequences: PDFHDK-*YGNAVLASGTAF*-CVATWV in COX7B2 versus PDFHDK-*YGNAVLASGATF*-CIVTWT in COX7B (six residues shown on either side). Given that the paralogous peptide is processed and presented, presentation of the COX7B2 peptide is plausible.
3. DMRTB1 — YYLPPPPPP. Not seen in IEDB.
4. TEX14 — EKDISLTDI. Not seen in IEDB.
5. RAD51AP2 — VYLKEIAKK. Not seen in IEDB.
6. PABPC3 — PAAAAAAAAT. Not reported in IEDB. We did not have a synthetic standard for this peptide.
7. RBM46 — SIEVTLAK. Not reported as such, but the N-terminally extended form ASIEVTLAK has been identified specific to colorectal cancer specimens (Yao et al., 2024). Because timsTOF instruments are prone to in-source fragmentation (Schramm et al., 2026), SIEVTLAK could be an in-source fragment of ASIEVTLAK.
8. EFCAB8 — MNLVAVASTRQKID. Not seen in IEDB.
9. OVOL3 — DAYIPGGPL. Not seen in IEDB.

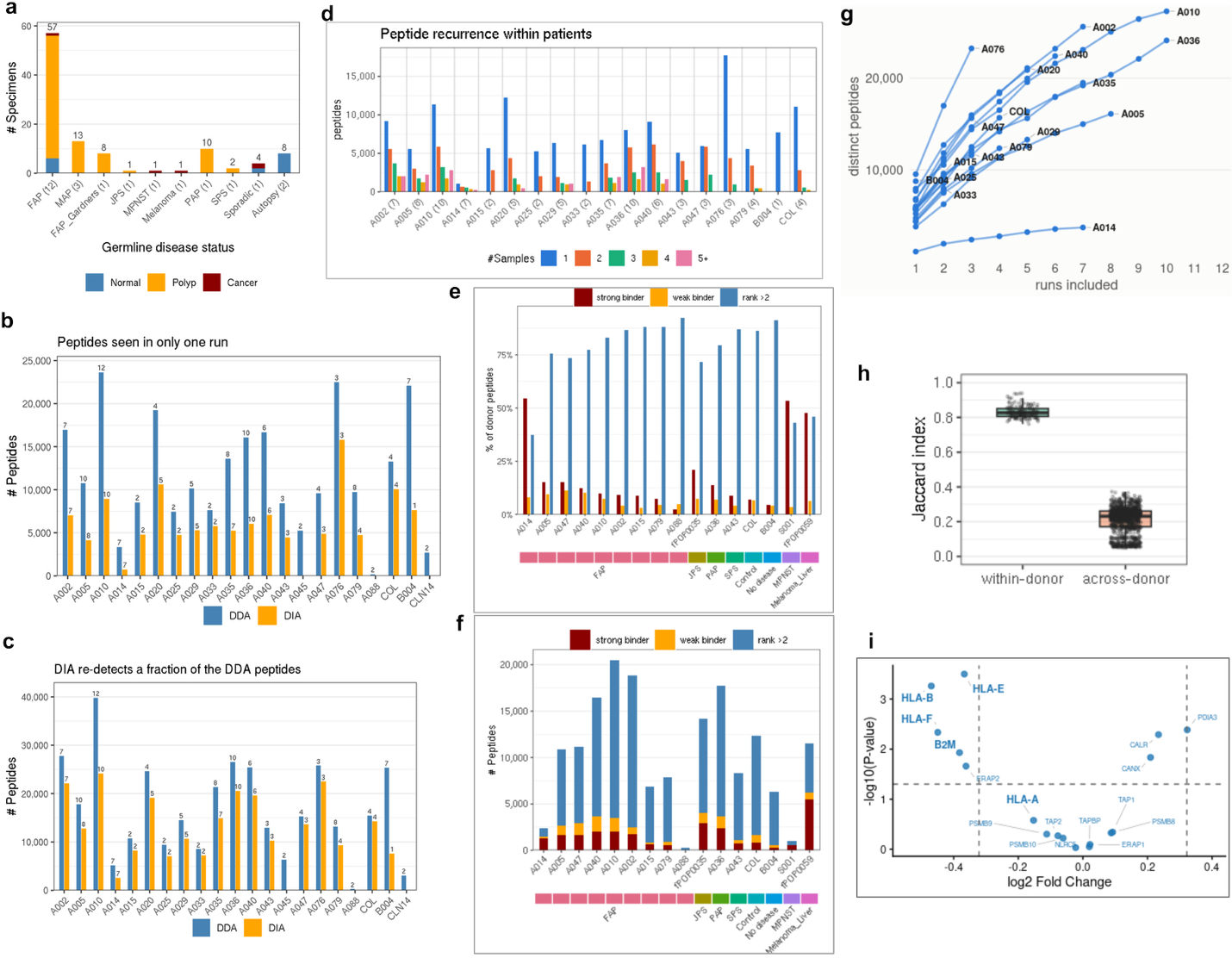

**Supplementary Figure 1**. Cohort composition, acquisition-mode comparison, and depth of the immunopeptidome dataset. **a)** Number of specimens profiled, grouped by germline disease status (number of donors per group in parentheses) and colored by tissue type. **b)** Peptides detected in only a single MS run per donor, shown separately for DDA (blue) and DIA (orange). Numbers above bars give the number of samples contributing to each acquisition mode. **c)** Total number of distinct peptides identified per donor by DDA (blue) and DIA (orange). **d)** Peptide recurrence within donors. Bars show the number of peptides detected, coloured by the number of samples in which each peptide was found. Sample count per donor is given in parentheses on the x-axis. **e)** Predicted HLA class I binding from NetMHCpan 4.1 of donor-specific peptides, shown as the percentage of each donor’s peptides classified as strong binders (%BA_rank or %EL_rank  ≤ 0.5, dark red), weak binders (0.5 < %rank ≤ 2, orange), or non-binders (%rank > 2, blue). The coloured bar beneath the x-axis denotes germline disease status. **f)** Same data as in (e), shown as absolute peptide counts per donor. **g)** Cumulative number of distinct peptides (log scale) as a function of the number of MS runs included, with one line per donor. Curves have not plateaued for most donors, indicating the immunopeptidome remains incompletely sampled at current depth. **h)** Volcano plot of HLA-I antigen-presentation machinery RNA (Esplin 2024) in polyp versus normal tissue. Dashed horizontal line, p = 0.05; dashed vertical lines, fold-change cutoff. HLA-I and β2M are reduced in polyps, while chaperones of the peptide-loading complex (PDIA3, CALR, CANX) are increased.

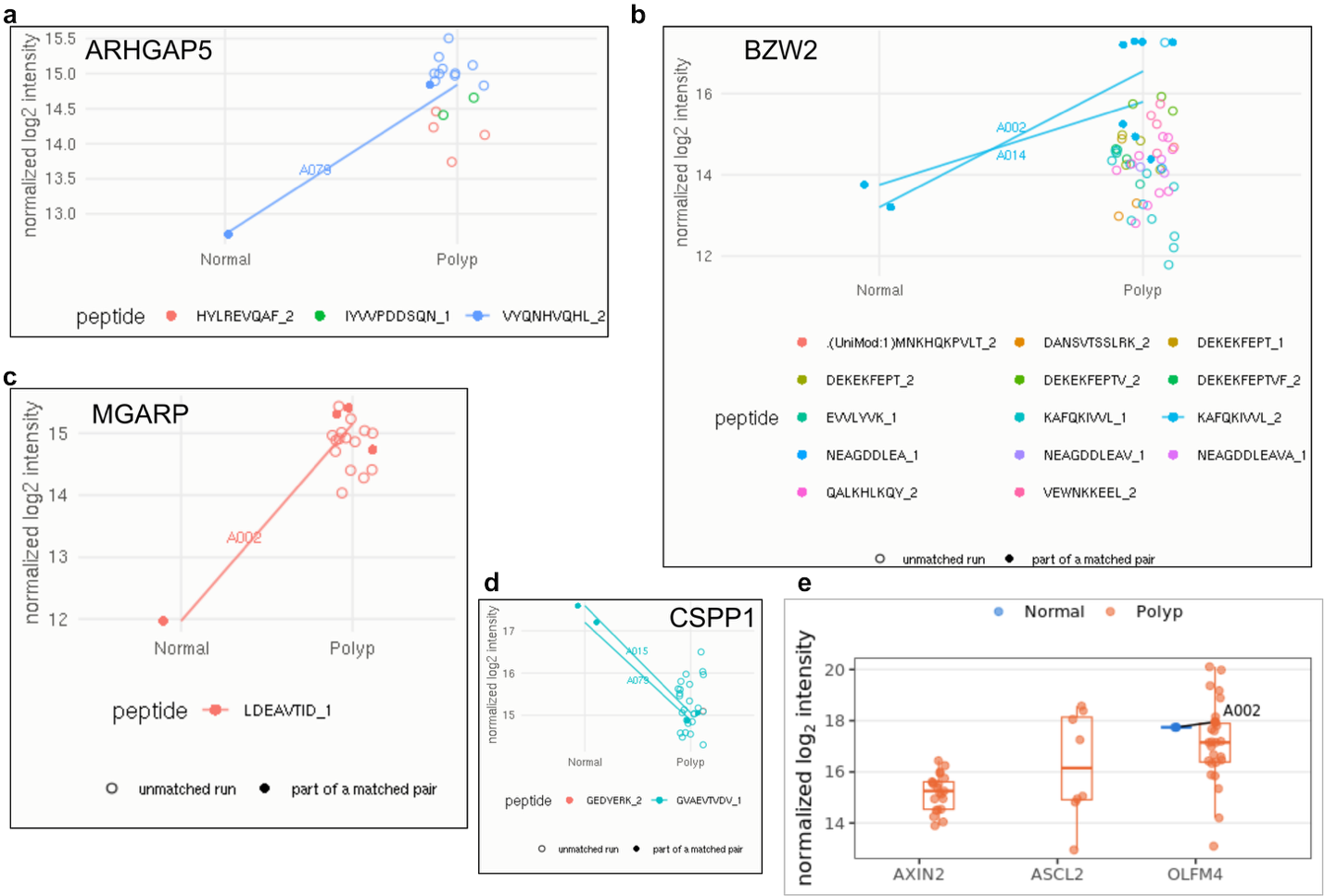

**Supplementary Figure 2**. dia-PASEF quantitation for all peptides of selected source genes. **a)** ARHGAP5, **b)** BZW2, **c)** MGARP and **d)** CSPP1, in normal versus polyp tissue. Each point is one precursor in one MS run, coloured by peptide sequence (charge state given after the underscore). Filled points joined by a line are measurements from the same donor with matched normal and polyp tissue, labelled by donor identifier; open points are runs without a matched counterpart. **e)** Normalized log2 intensity of peptides derived from Wnt/β-catenin target genes (AXIN2, ASCL2, OLFM4) in normal and polyp tissue. One point per MS run; solid points joined by a line are per-patient means for patients with more than one tissue sampled. Boxes show median and interquartile range, whiskers extend to 1.5× IQR.

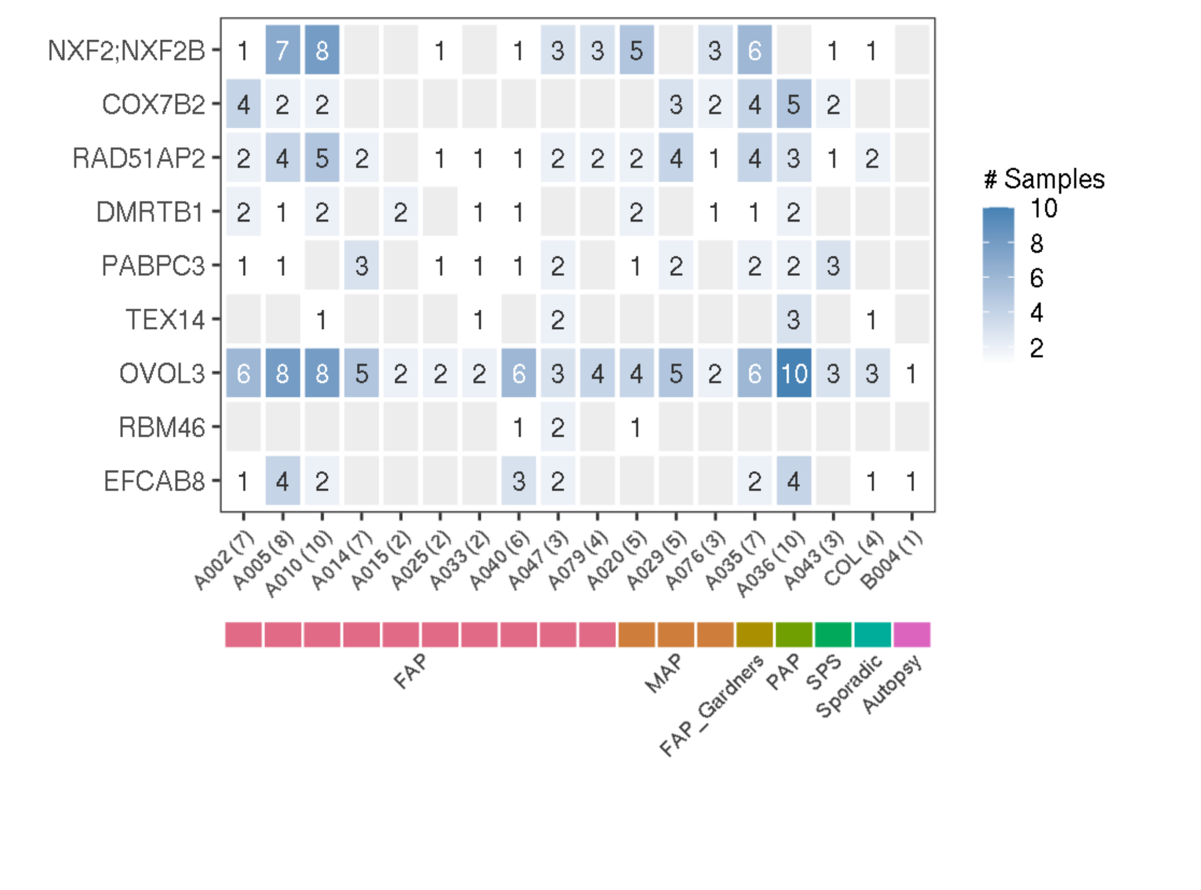

**Supplementary Figure 3**. Heatmap showing, for each candidate antigen source gene, the number of samples per donor in which the peptide from that gene was identified in the immunopeptidome; grey indicates no detection. Patient identifiers on the x-axis include the number of samples per patient in parentheses, and the coloured bar beneath denotes germline disease status.

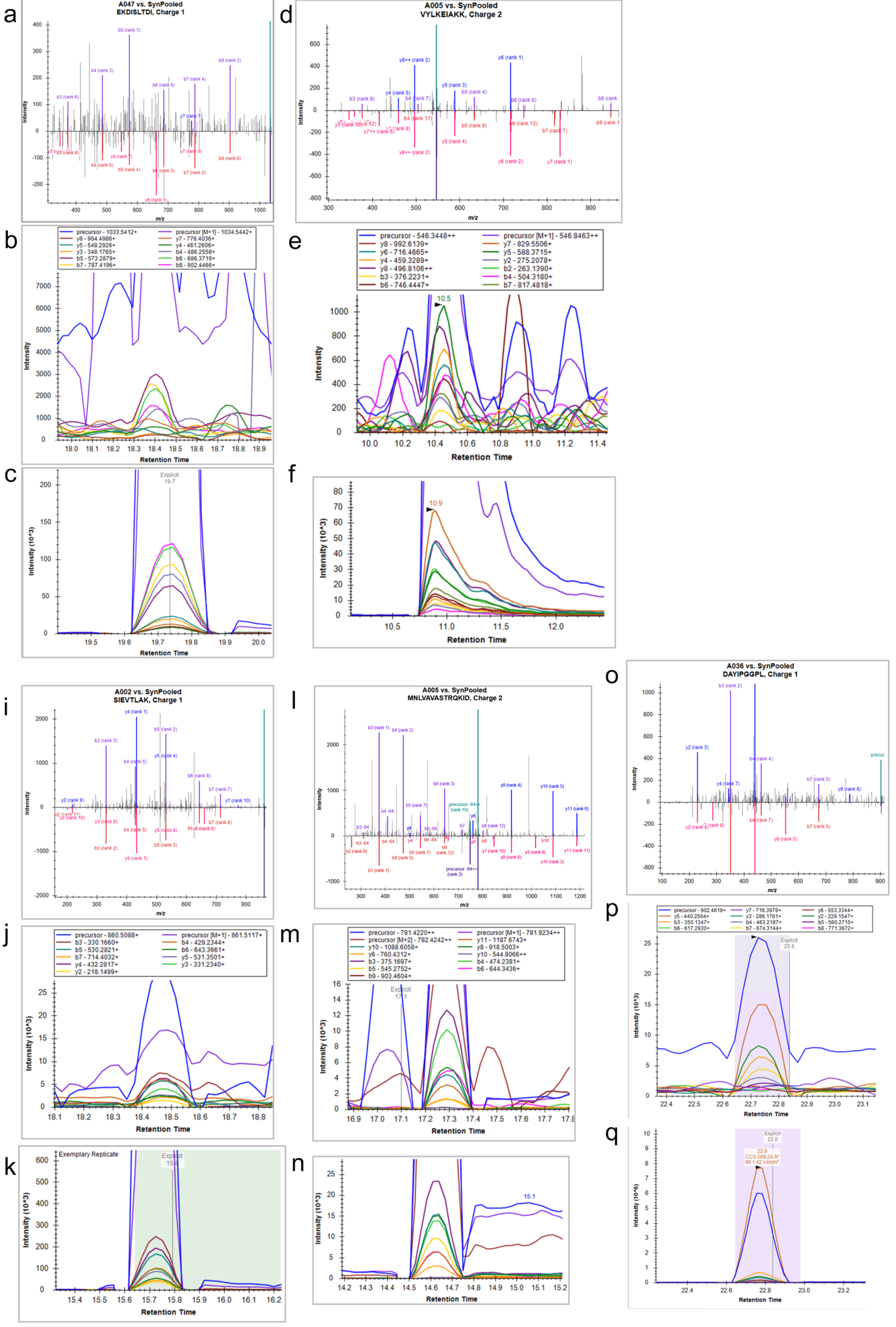

**Supplementary Figure 4**: **a)** Mirror plot of the dda-PASEF spectrum for TEX14 antigen EKDISLTDI(+1) from donor A047 (polyp, Sample ID 97) against the corresponding synthetic peptide standard. **b)** XICs from donor A047 (polyp , Sample ID 69) for the same peptide as in (a). **c)** XICs from synthetic peptide standard for the same peptide as in (a).

**d)** Mirror plot of the dda-PASEF spectrum for RAD51AP2 antigen VYLKEIAKK(+2) from donor A005 (polyp, Sample ID 55) against the corresponding synthetic peptide standard. **e)** XICs from donor A005 (polyp , Sample ID 55) for the same peptide as in (d). **f)** XICs from synthetic peptide standard for the same peptide as in (d).

**i)** Mirror plot of the dda-PASEF spectrum for RBM46 antigen SIEVTLAK(+1) from donor A002 (polyp, Sample ID 94) against the corresponding synthetic peptide standard. **j)** XICs from donor A047 (polyp , Sample ID 69) for the same peptide as in (i). **k)** XICs from synthetic peptide standard for the same peptide as in (i).

**l)** Mirror plot of the dda-PASEF spectrum for EFCAB8 antigen MNLVAVASTRQKID(+2) from donor A005 (polyp, Sample ID 24) against the corresponding synthetic peptide standard. **b)** XICs from donor A047 (polyp , Sample ID 97) for the same peptide as in (l). **c)** XICs from synthetic peptide standard for the same peptide as in (l).

**o)** Mirror plot of the dda-PASEF spectrum for OVOL3 antigen DAYIPGGPL(+1) from donor A036 (polyp, Sample ID 72) against the corresponding synthetic peptide standard. **p)** XICs from donor A036 (polyp , Sample ID 72) for the same peptide as in (o). **q)** XICs from synthetic peptide standard for the same peptide as in (o).

XIC: Extracted - ion chromatogram

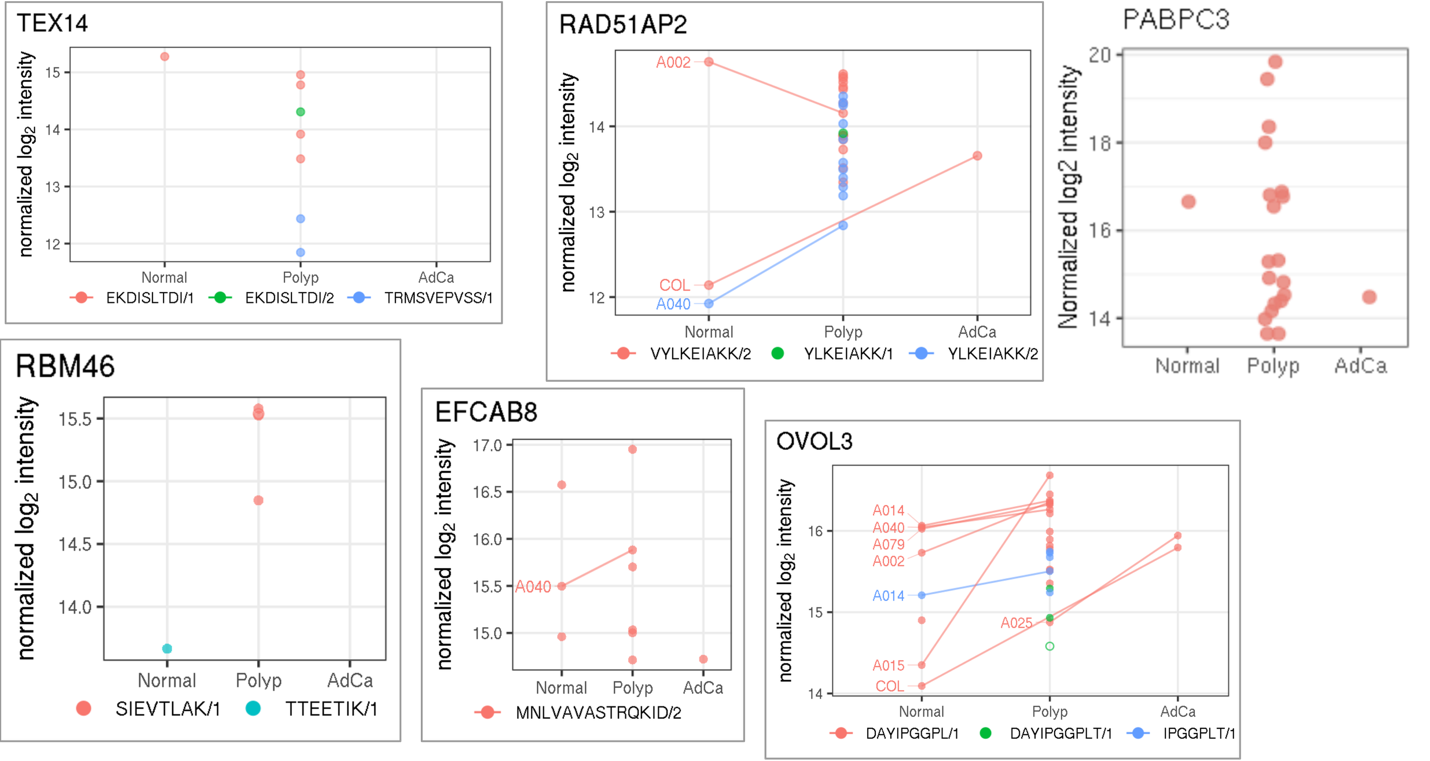

**Supplementary Figure 5**: Normalised log2 intensity of six candidate peptides from source genes TEX14, RAD51AP2, PABPC3, RBM46, EFCAB8, OVOL3 (A0AA75L0V4) across disease stages, coloured by precursor charge state. Specimens with matched mucosa are connected with lines.

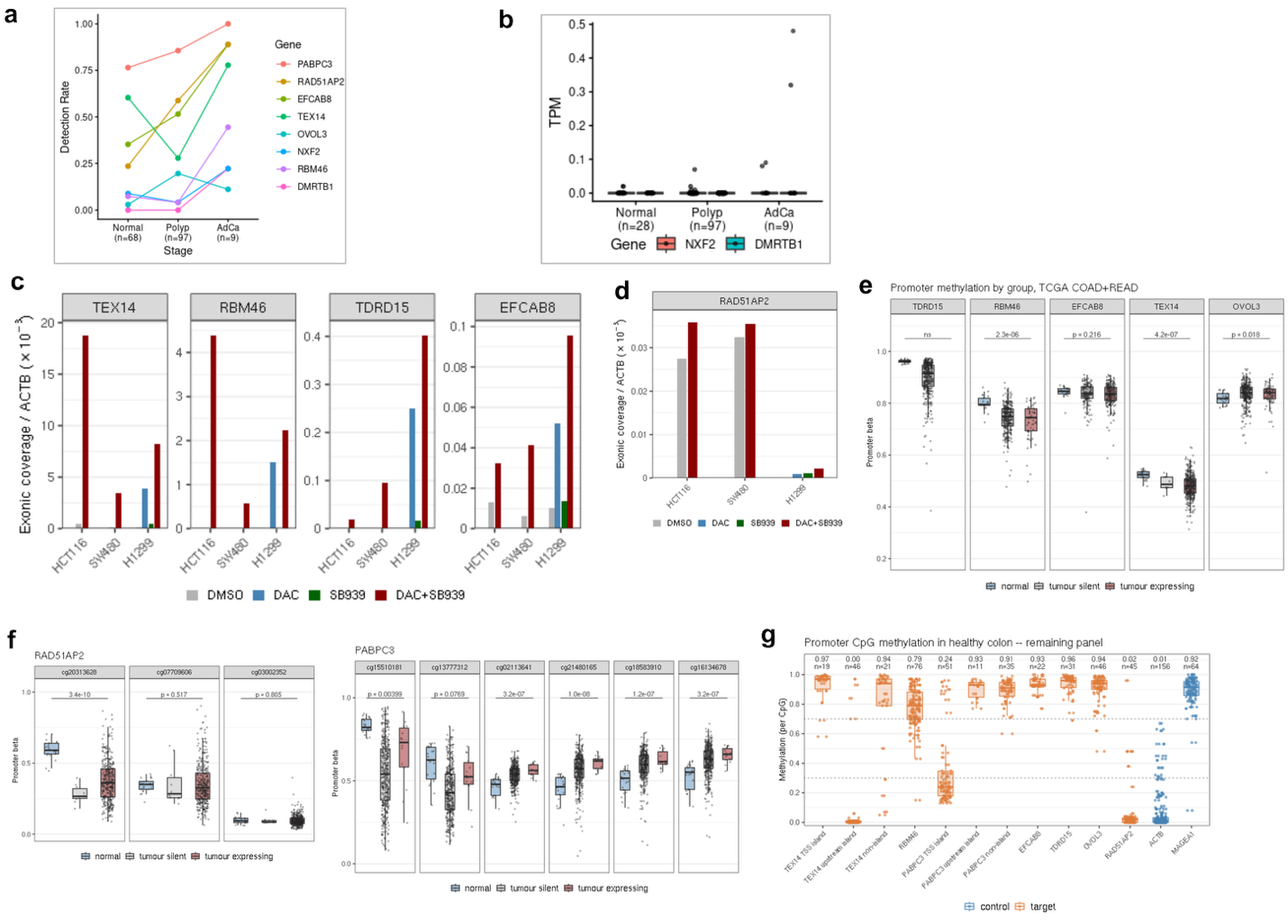

**Supplementary Figure 6**. **a)** Detection rate, calculated as the number of samples with expected count > 0 divided by the total number of samples. **b)** STAR TPM in FAP polyps for NXF2 and DMRTB1. **c)** Exonic coverage of TEX14, RBM46, and EFCAB8 normalised to ACTB. **d)** Exonic coverage of RAD51AP2 normalised to ACTB. **e)** Promoter CpG methylation (Illumina 450K β values) for RBM46, EFCAB8, TEX14 and OVOL3 in TCGA COAD and READ, split into normal tissue, tumours not expressing the gene, and tumours expressing the gene. p-values from unpaired Wilcoxon rank-sum test. **f)** As in (e), shown per probe. Left, RAD51AP2. Right, PABPC3. **g)** Per-CpG promoter methylation in healthy colon (Roadmap E106 WGBS) for TEX14 loci, RBM46, PABPC3 loci, EFCAB8, OVOL3, RAD51AP2, and for the controls ACTB and MAGEA1. Numbers above each group give the median methylation and the number of CpGs assessed.

DAC: Decitabine; SB939: histone deacetylase inhibitor pracinostat.

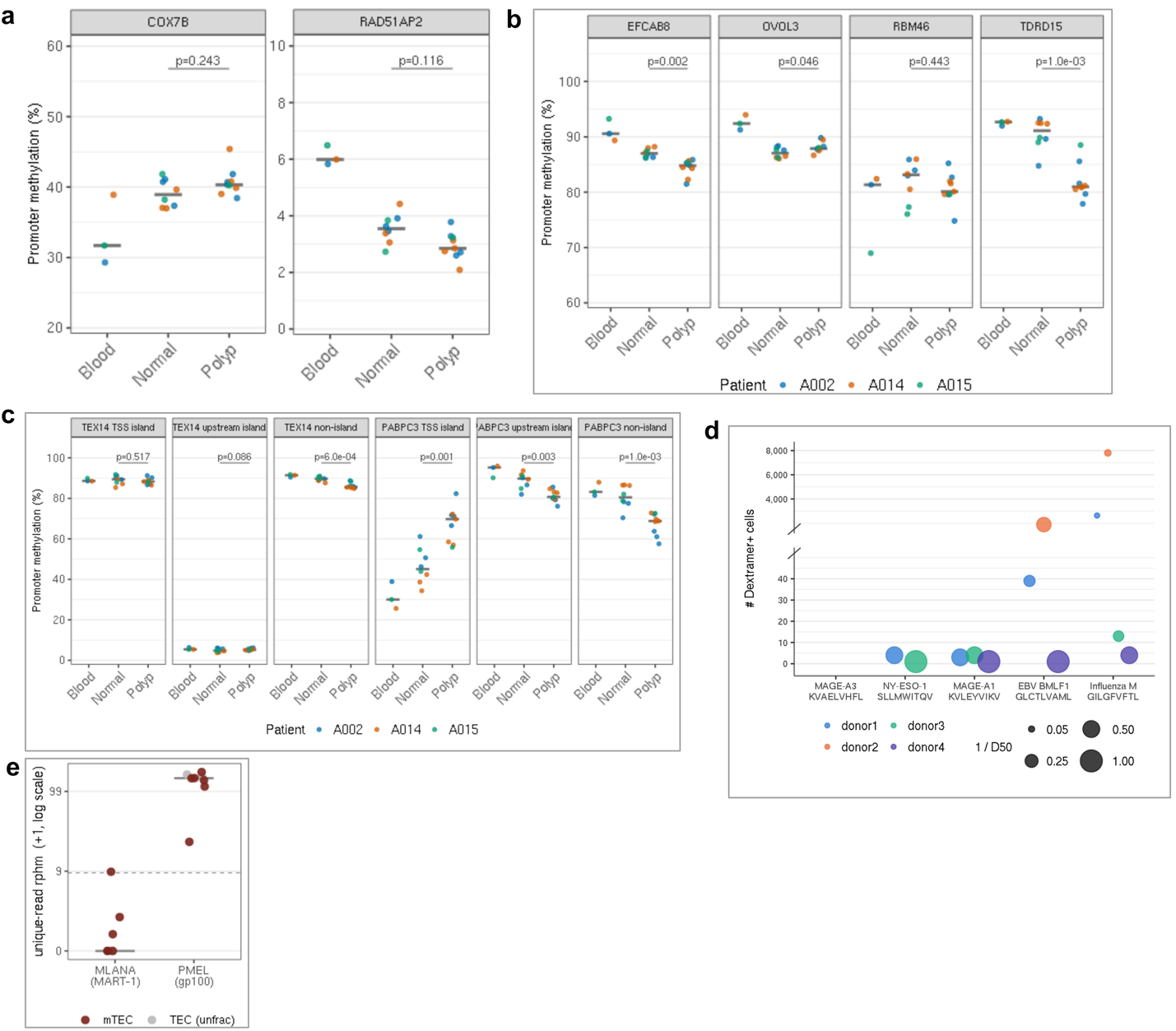

**Supplementary Figure 7**. **a)** Promoter CpG methylation for COX7B, and RAD51AP2 in blood, matched mucosa and polyp from three FAP donors. Each point is one specimen; horizontal bars show the group median. p-values from two-sided within-donor permutation test. **b)** As in (a) for EFCAB8, OVOL3, and RBM46. **c)** As in (a), split by promoter region for TEX14 and PABPC3. **d)** TCRβ repertoire of peptide-dextramer-sorted T cells against reference CTA and viral epitopes, from the 10x dataset. Each point is one donor; the y-axis gives the number of dextramer+ cells, and point size gives 1/D50, where D50 is the fraction of clonotypes accounting for 50% of reads. Larger points indicate more clonally focused repertoires. Peptides are labelled with their source gene and HLA-I allele of dextramer used (Supplementary Table5). **e)** Expression of the MART-1 and gp100 genes in medullary thymic epithelial cells (mTECs) and thymic epithelial cells (TECs). Each point is one donor; horizontal bars show the median. Dashed line marks the 8.55 rphm detection threshold.
